# The speed limits of memory

**DOI:** 10.1101/2025.03.31.646314

**Authors:** Bastian Wiederhold

## Abstract

As anyone who has tried to memorize a one-hundred-digit number can attest, acquisition of numerical information typically proceeds at less than one bit/s. If human memory operated at this speed in general, even a simple conversation would not be possible. Indeed, through techniques such as the memory palace, which translate numerical information into more natural contexts, memory athletes manage considerably higher rates. This suggests that memory, in its intended environment and under training, performs at substantial speed, which can be quantified in memory competitions. Analyzing the data reveals three phenomena. First, in short-duration tasks up to 42 bit/s have been achieved. Remarkably, competitors spend most of the time on reading, indicating that they form mental associations even more rapidly. Second, record performances show a remarkable concordance across time scales: the processing speed depends on memorization time as a power law. Third, despite dramatic improvements in scores and mnemonic strategies over the last decades, the differences in information rates across memorization tasks remain remarkably consistent.

---

Memory techniques have been known since antiquity^1^. Starting about three decades ago, formal rules for competitions were established, which allowed a comparison of the competitors’ control of mnemonic strategies. Nowadays, across different disciplines such as numbers, cards, word, faces and images, common scores at competitions are several times higher than expected from naive memorization. The strategies can be interpreted as converting abstract information into more natural and familiar structures^2^. An example is the famous “ memory palace” or “ method of loci”, which is the dominant approach for several disciplines. During memorization, competitors first transform information into “ mnemonic images”: memorable objects, people or scenes; more natural entities. These images are then linked to prominent locations along a mental walk; a more familiar sequential structure. During recall, competitors reimagine the same mental journey and decode the mnemonic images to retrieve the stored information. Viewing the techniques as ways to obtain more natural and familiar representations of information, the competitions might inform us about the true speed limits of memorization.

Recently, it has been argued that the human brain is “slow”, processing information only at around 10 bits/second including examples from memory competitions^3^. Our systematic analysis will reveal a more diverse picture. We analyze data from classical competitions of the International Association of Memory (IAM), the modern format Memory League (ML), and speed-memory.com. Combined, these tasks span a wide range of memorization-time periods from one second to one hour. To compare different disciplines, we calculate the minimum information processing rates in bit/s necessary to explain top performances. We find that for short tasks, information rates greatly exceed 10 bit/s and seem to be limited by reading rather than memorizing. Quite generally, the rates follow a power law as a function of the memorization time.

## Memorization at nearly the speed of reading in short tasks

Using a memory palace involves three processes: reading, associating and navigating (Figure 1 **A**). For instance, competitors first map visual input in form of digits onto specific prememorized mnemonic images. This process is comparable to standard reading as, indeed, the most popular strategy to learn the map is to assign letters to the digits 0 − 9^4^. The resulting images are associated to salient positions of the mental walk through short creative stories. Once a location is occupied, the competitor mentally navigates to the next location of the memory palace. Among these three processes, it seems most natural to equate “ memorization” with the process of forming associations.

**Figure 1.**
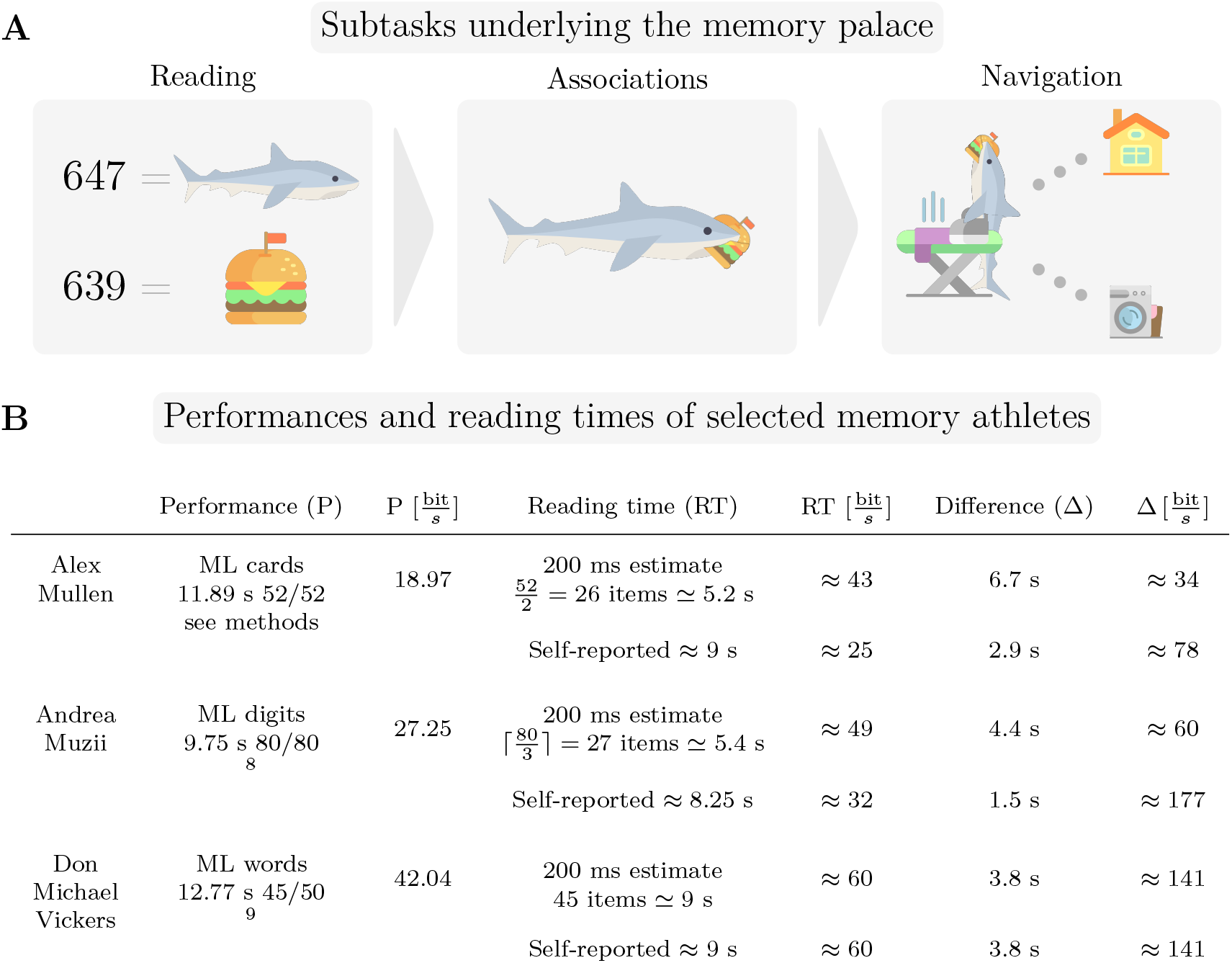
**A** Schematic of the process of using a memory palace. First, the input, in this case numbers, are transformed into mnemonic images. Second, associations are formed between the mnemonic images and also the position of the mental walk. Once a position is occupied, competitors navigate mentally to the next salient location. In most cases, the mental walk is along a real-world location. **B** Performances, estimated minimal and self-reported reading times, differences Δ between the reading and memorization time and the associated information rates for selected memory athletes. To determine the length of the reading phase, we asked the competitors and estimated the minimum based on 200 ms per item^5^. Even the latter conservative estimates constitute a large fraction of the memorization time. If reading and the formation of associations were entirely sequential, Δ would be the association/memorization time corresponding to the speed in the right-most column. In ML cards the task is to memorize the order of 52 cards in a shuffled deck, in ML digits an 80-digit decimal number and in ML words a sequence of 50 words. Andrea Muzii is using a three-digit system, converting three-digit decimal numbers into one mnemonic image, so there are 27 items to be perceived^10^. Alex Mullen achieved the time with a system encoding two cards in one mnemonic image, so 26 items need to be perceived^11^.

To understand the time spent on the three processes, we asked top competitors in ML cards, digits and words for the time they require to read the items of the tasks (Figure 1 **B**). The self-reported reading times could be fairly accurate, as competitors train extensively over years to minimize total time. In ML words, reading is comparable to everyone’s understanding of the term, whereas in ML cards and digits, reading refers to the aforementioned transformation into mnemonic images.

Furthermore, we calculated a realistic minimum of the reading time based on research on single-word reading, which suggests that 200 milliseconds elapse due to visual processing before the visual input becomes available to the mnemonists’ minds^5^. We received values in line with the self-reported times and the estimate of 50 bit/s for general reading^6^.

Self-reported times and the estimated minima indicate that reading requires most of the memorization time. If mental associations were formed solely during the difference Δ between the memorization time and the reading time, processing rates in memory would be significantly higher than indicated by the task itself.

### Power law of memorization speed

Across memorization tasks, the greater the number of items to be memorized, the longer the time competitors require. The reading time per item is not expected to increase decisively, but more time is required to consolidate the information in memory until the start of the recall phase. It is, therefore, not surprising that the information rates are lower for longer tasks. Intriguingly, when we calculated the minimum information rates necessary to explain the records, we found a power law of information rate over the memorization time (Figure 2 **A**). We fitted the function *aT* ^*b*^ for two parameters *a, b* to the data points provided by the official records using least squares based on log quantities. As a function of the duration *T* of the memorization phase in seconds, the resulting curve of the information rate *R* is

**Figure 2.**
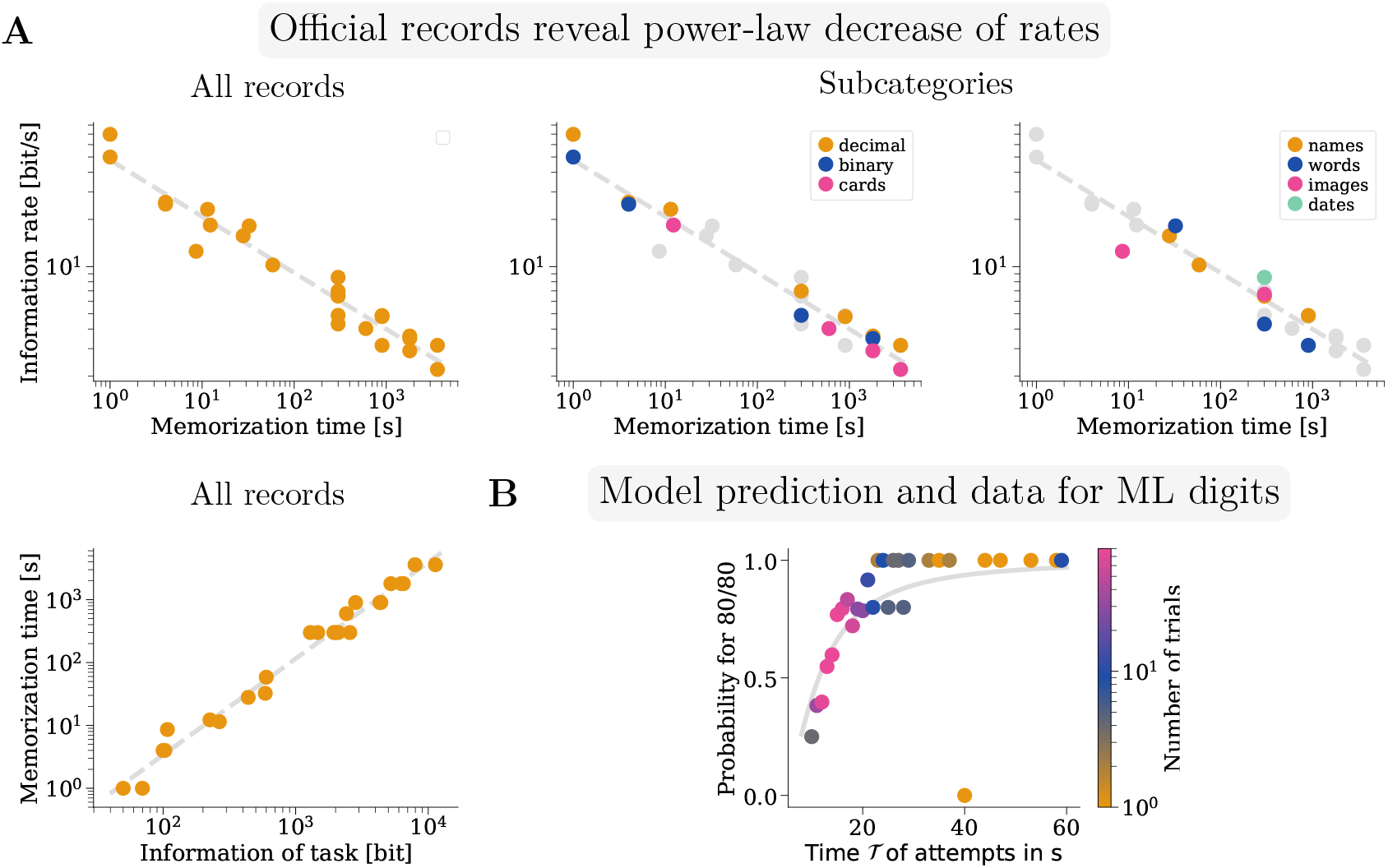
**A** The left panel in the top row depicts the information rates of all official memory world records as a function of the duration of the memorization time. The results of 1 s and 4 s memorization duration are from speedmemory.com. The remaining results with a memorization time smaller than 60 s are ML records and the bottom right-hand cloud corresponds to the records from classical competitions. The vertical alignment of parts of the data results from the memorization time being either 5, 15, 30 or 60 min at classical competitions. The speed-card record in classical competitions of Shijir-Erdene Bath-Enkh is 12.74 s. It is reassuring that the score is similar to the ML cards record of 12.25 s, as this is the only discipline directly comparable between ML and classical competitions. The middle and right panel of the top row show the same records, but grouped by discipline categories. The bottom panel shows instead memorization duration as a function of the information of the complete tasks. **B** Empirical success probabilities for achieving 80*/*80 in ML digits compared to the prediction of the probabilistic model. Shown are data for the success probability of trials, binned in intervals of one second, by competitors Alex Mullen and Andrea Muzii, the official and unofficial ML record holders. Note that around 85% of trials are in the time between 10 and 20 s, and larger times are insufficiently sampled. For example, there was only a single attempt in the 40 s bin. The curve depicts the prediction ℙ (*A*(80, *T −r*)) of the model. We used the parameters of the power law 1, a reading time *r* of 8 s (Figure 1 **B**) and obtained *p* = 0.72 for the free parameter of the model using weighted least squares, with the number of attempts of a certain time as weights.

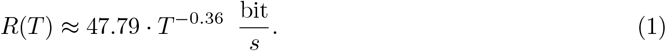

Note that, compared to standard forgetting power laws^7^, Equation (1) does not describe the proportion of correctly recalled items over time, but rather the achieved information rates over different time spans. The dependence of the memorization time on the information of the complete task is also described by a power law rather than a linear relationship (Figure 2 **B**). As a function of the complete memorized information *I* in bit, the memorization time in seconds is described by

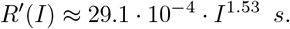

Using a simple probabilistic model, we related the power law to the probability to remain error-free in tasks with a certain number of items. The model is based on the assumption that competitors aim for error-free memorization and that long tasks can be divided into shorter tasks with independent error probabilities. The model predicts that, given enough time, competitors’ success-rate for perfect memorization of a certain number of items is high, but rapidly decreases if the memorization time becomes shorter than a certain threshold. This seems to match observations on the error-rates in ML (Figure 2).

### Consistent differences between cards, decimal and binary digits

Classical competitions feature three symbolic coding systems: cards, decimal and binary digits. Over recent decades, world records have improved steadily with no clear indication that the trend is weakening (Figure 3 **A**). Among the disciplines based on symbolic sets, decimal digits stand out with strikingly linear record progression (Figure 3 **A**). Despite this development, there are consistent relations between the achieved information rates across symbolic sets. Decimal digits are processed faster than cards: this trend is convincing for disciplines from 10 s to 60 min memorization time over the past 30 years (Figure 3 **B**). The comparison between decimal and binary digits is more complicated: there is a clear and consistent gap of around 40% for the 1 s and 5 min disciplines, but essentially the same memorization speed in the 30 min discipline (Figure 3 **B**).

**Figure 3.**
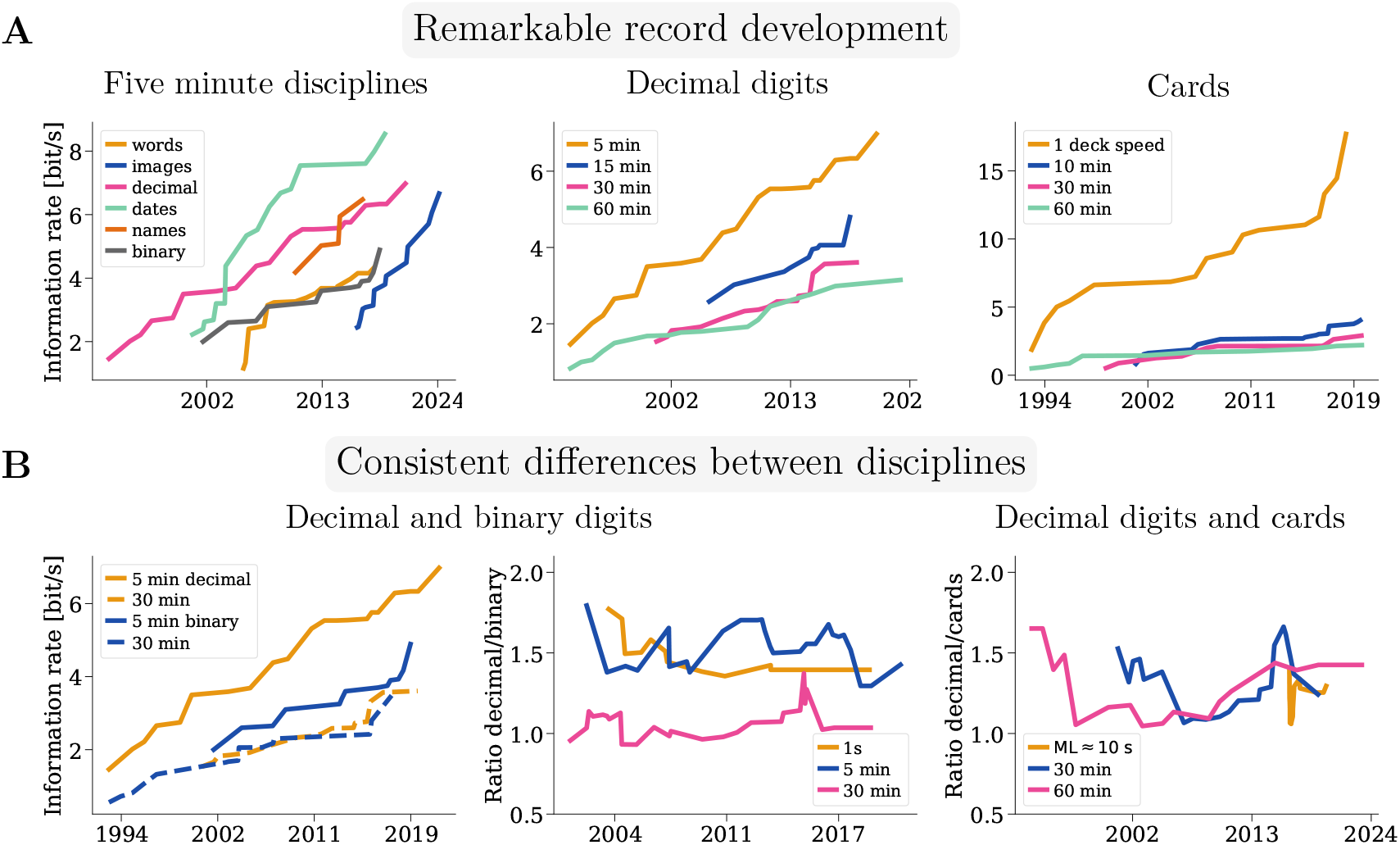
**A** Record development of selected disciplines since the inception of memory competitions. Left, all disciplines with 5 min memorization time. Middle, classical disciplines based on memorizing long sequences of decimal digits. Right, classical disciplines based on playing-card memorization. **B** Development of the information rate differences between decimal and binary digits, as well as cards. Left, record development of 5 and 30 min decimal- and binary-digits memorization. Middle, ratio of the information rates of all decimal and binary digits disciplines of comparable length. Right, ratio of the information rates of all decimal-digits and cards disciplines of comparable length.

## Discussion

A recent study revealed the conundrum of human cognition: whereas our sensory systems are able to process enormous quantities of information, in the range of 10^9^ bit/s for the retina, most human activities, from playing Tetris over memorization to typing, exhibit only about 10 bit/s of processing^3^.

The prevalence of parallel processing in sensory systems and serial processing in high-level cognition was identified as one of the major contributing factors. The significant fraction of time spent on reading in rapid memorization tasks, performed at up to 42 bit/s, illustrates that either there is more high-level parallelization than suggested or that, excluding perceptual time, the human brain can be way faster. If reading, associating and navigating were parallel, this would illustrate that even though humans seem to perform one action at the time, highest human processing speeds can be supported by an underlying parallelization of the involved brain regions. If the processes were instead sequential, the narrow difference between memorization and reading time would imply that the actual process of memorization reaches around 150 bit/s. Equally, any combination of the two options would explain part of the conundrum. It would be interesting to understand the speed of different cognitive subprocesses and how much parallelization exists during the use of memory palaces by top competitors.

The power-law decrease in the bit rate with time across multifarious representations of information (numbers, cards, words, faces, images) is a sign that the analysis reveals a general principle about the rate at which associative systems in the brain store information. It is plausible that the power-law is a consequence of a larger time investment in revision and in better and more vivid associations to overcome the forgetting curve^12,13^. The probabilistic model suggests that the power law relates to the diminishing returns of the success probability of competitors to memorize certain information as time increases. During memorization, competitors may read the information only once at a slower pace, or revise twice or even several times in longer disciplines. As, additionally, the number of attempts on longer disciplines is smaller and we have no statistics on competitors errors, future controlled experiments will uncover the precise principles of the power law.

The consistent gaps in processing speed suggest that the difficulty of encoding systems for human beings goes beyond the amount of conveyed information. The 52 options in playing cards add a complexity which is not present when memorizing decimal digits.

The 40% gap between binary and decimal digits in the one-second and five-minute disciplines indicates that decimal digits strike a good balance between readability and complexity. We suppose that three factors explain the difference. First, the effect of word-length as more symbols are necessary to transfer the same information^14^. For instance, more saccades might be necessary to accumulate the information or more peripheral vision is used, which could explain lower information rates^15^. Second, the effect of reduced salience as visual binary information is more redundant than decimal information^16^. Since binary saccade targets are less prominent, competitors might need to correct initial eye movements more often. Both effects can be experienced when reading the numbers

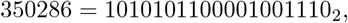

although the newest competition software tries to overcome the limitations through horizontal separation bars and colored cursor highlights^17^. Third, competitors use systems for binary digits, which either carry less information per mnemonic image or were trained less. The most straightforward way to memorize binary digits is to convert three-digit binaries, say 010, into a single decimal digit, here 2. There are eight three-digit binary numbers, so that the strategy leads to a decrease of memorized information by

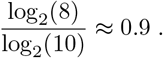

Competitors start by learning a system to memorize decimal digits, as decimal information is more ubiquitous in real life and featured in other disciplines in speed-memory.com and classical competitions. Binary-specific systems, overcoming the above factor, might not be trained at all or not to the same level of proficiency.

The assimilation of processing speeds in the 30-min disciplines might be linked to the grading systems: decimal digits are presented in rows of 40 (log_2_(10^40^) ≈133 bits), whereas binary digits are provided in rows of 30 (log_2_(2^30^) = 30 bits). If the row is correct 40 respectively 30 points are awarded. If a single digit is different, points are halved and no points are awarded if more than two digits are incorrect. As practically all mnemonic systems for decimal and binary digits encode at least two digits with an mnemonic image^2^, an error results in the loss of all points for the row. Thus, the grading system punishes an error in decimal digits with a deduction of 133 bits and in binary digits only with 30 bits. In the half-an-hour discipline forgetting and errors might become more important than the ability to parse information. This might force competitors to revise or spend additional time for decimal compared to binary digits.

The information rates we calculated represent lower bounds on the speed of processing. In classical competitions records may contain errors, which lead to significant deductions in points. For associative disciplines such as names and faces or historic dates, we used a simplification that considers the associative elements to be as clearly distinguishable as possible. We did not model human memorization of written historic events, images or faces, which likely depends on manifold sentence/image properties^18^ and domain expertise^19,20^. Equally, the use of memory palaces requires additional visuospatial and contextual information to be processed, which we did not consider. In general, the rules of memory competitions, such as grading schemes, are not precisely aimed at maximum information processing by competitors.

In conclusion, our systematic analysis reveals that the speed of memory is insufficiently described by assuming a universal limit of 10 bit/s. The significant differences between encoding systems illustrate that human beings are influenced by many aspects which go beyond the pure information-theoretic content of a task. Over a short period, human beings are able to memorize at around 50 bit/s, but the rate decays with longer memorization times according to a power law. A major fraction of the required time in short tasks can be explained by reading. This indicates that, if one excludes perceptual time, the formation of associations in memory might even occur at a rate of more than 100 bit/s.

## Acknowledgements

I would like to thank Markus Meister, who drew my attention to the information rates of human performances.

This paper would not have been possible without the support of the memory sport community. I am grateful to Johannes Mallow and Simon Orton for sharing the dataset used to comment on past performances of top competitors during ML events. Johannes Mallow helped with several questions. Simon Orton kindly provided information on the generation of words and names in ML. I am thankful to Katie Kermode for discussions and for providing me with information on the generation of names and words in the classical competitions. Alex Mullen, Andrea Muzii, Don Michael Vickers and Enrico Marraffa generously gave insight into their performances.

Jan Rapp pointed out improvements of information calculations. Andreas Herz and Martin Stemmler contributed with stimulating discussions and helpful feedback on the manuscript.

## Methods

### Data acquisition

To analyze classical competitions, we used the world record histories^21^ from the statistics website^22^ of the International Association of Memory (IAM), where also the official online world records for ML can be found^23^. Results are only counted as world records if the result is obtained during an online competition with videos recorded from two different angles. Separate in-person records are kept but are outdated as in-person competitions have ceased to exist for ML since the COVID-19 pandemic. The world records and development for the 1 s and 4 s decimal and binary disciplines can be found on speed-memory.com^24,25^.

Johannes Mallow (commentator on ML competitions^26^ and former world champion) and Simon Orton (ML developer) provided us with a database of 191503 results of top competitors from the recent results on the ML website^27^ covering the period between June 2017 and January 2025. The dataset is public as the main page of the ML website allows to search recent results arbitrarily far back. Success statistics, as the one depicted in Figure 2 **B**, are frequently shown to assess the likelihood of participants to get a certain score during online competitions. We generated unofficial ML world records, including the ML cards performance of 11.89 s by Alex Mullen, by searching the database for the best results. A list of all records is at the end of the document.

### Competitions and disciplines

A companion work elaborates on the used mnemonic techniques and provides additional background information on the competitions^2^.

### Classical format

Memory competitions started with the first world championship in 1991, which initiated the classical format: a mental decathlon featuring the memorization of decimal, binary and auditive digits, cards, words, images, names and historic dates (Table 1).

**Table 1:** Overview of the disciplines in classical competitions. The first seven disciplines involve the memorization of long sequences. Decimal and binary digits, cards and words are presented in rows of *k* items: a single error in a row leads to the row being counted as *k/*2, whereas with two or more errors no points are being awarded. Speed cards is the only discipline in which the time rather then the amount counts. The number of cards counts until the first error - however only an error-free performance leads to a competitive score. In images recall competitors need to indicate the previous permutation of the five images in a row with a deduction of one point, if an error is made. As images rows contain only five items, strategies which do not involve knowing the overall sequence, but only the sequence per row, are possible. Historic dates and names and faces are associative disciplines. In recall, the events or the faces are in a random permutation and competitors need to reconstruct the year respectively the name. The disciplines differ in our ability to estimate the information rate processed by human beings in bit/s. For the purely number or card based disciplines entropies are straightforward to calculate. For images and historic dates we use theoretic lower-bounds, whereas for word and names we employ estimates based on database samples.

|  | memorization objective | memorization time [min] | grading scheme |
| --- | --- | --- | --- |
| decimal digits | a sequence of decimal digits | 5, 15, 30, 60 | rows of 40 |
| binary digits | a sequence of binary digits | 5, 30 | rows of 30 |
| auditory digits | a sequence of decimal digits, read aloud at one digit/s | up to 8 | counts until first error |
| cards | a sequence of playing cards in the form of successive scrambled decks of 52 playing cards | 10, 30, 60 | rows (decks) |
| speed cards | the sequence of one scrambled deck of 52 playing cards as quickly as possible | at most 5 | essentially perfect |
| words | a sequence of words | 5, 15 | rows of 20 |
| images | a sequence of images with arbitrary motives in various art-styles | 5 | rows of 5 |
| historic dates | associate years between 1000 and 2100 to fictional events | 5 | deduction (guessing not allowed) |
| names and faces | associate international names to faces (first + last name) | 5, 15 | no deduction (guessing allowed) |

There are three versions of the events taking place over the course of one, two or three days with correspondingly longer disciplines. The three-day standard is reserved for the world championships. The memorization times of the disciplines range from 5 to 60 minutes with recall twice or thrice as long. For example, all standards feature 5 min decimal digits, and 15, 30 and 60 min decimal digits respectively in the one, two and three-day format. To determine the overall winner, the raw scores of the disciplines are converted to a point scale with a system inspired by the track-and-field decathlon. The precise rules can be found in the rulebook^28^ of the International Association of Memory (IAM).

### Memory League

In 2015 Memory League (ML) was launched, first under the name Extreme Memory Tournament^27^. There are fewer disciplines with much shorter memorization time, the focus is on one-vs.-one matches and top competitors optimize the time for a certain number of items, rather than optimizing the number of items per time. ML features the six disciplines decimal digits, cards, words, national names, international names, images (Table 2). The memorization time lasts up to one minute and recall up to four. Tournaments are based on set-based matches. During a set, players take turns choosing the disciplines, so typically all are featured in a set. Whoever made fewer errors wins the discipline and, if both have made the same number of errors, the person with the faster memorization time wins.

**Table 2:**
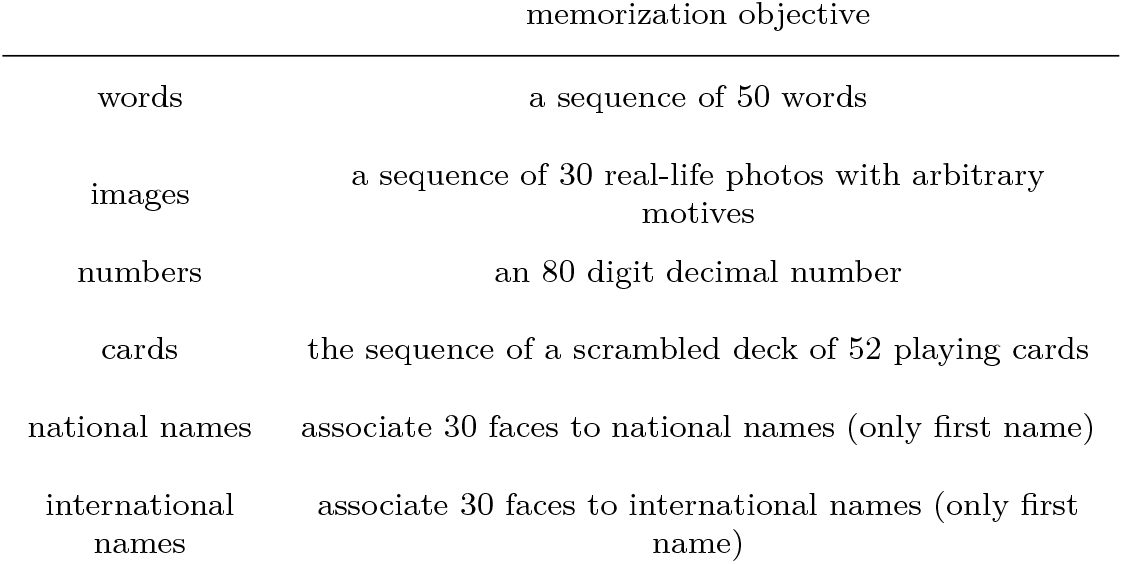
Overview of the disciplines in Memory League (ML). Compared to classical competitions the memorization time is shorter: at most one minute compared to at least five minutes in classical competitions with the exception of speed cards. Instead of optimizing the amount of information in a fixed time, competitors try to memorize a fixed-size set of information as quickly as possible.

### Speed-memory.com

The speed-memory.com competitions consist of six disciplines: 1 s and 4 s decimal and binary digits and the matrices and colored shape disciplines. The website^24^ is in Spanish and the competitive scene centered around Ramón Campayo and his students. At the time this article was written, it was unclear from the website whether there still was a competitive scene as there were no upcoming events and several sections of the website seemed outdated. In the 1 s and 4 s disciplines competitors see all information at once, but are allowed to place the decimal and binary digits in particular groups arranged across the screen.

### Information content of the different disciplines

Our calculations are based on Shannon’s entropy, defined as

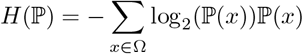

for a probability distribution ℙ on a discrete underlying event space Ω. Marking rules and the high number of possibilities prevent guessing of competitors. Therefore, if we calculate the entropy associated to the recalled items, it will correspond to transmitted information. Ideally, we would prefer to have a distribution of answers of a competitor. The records present only one realization, but at least the disciplines consist out of a sequence of repetitive microtasks such as individual digits. For all disciplines, our calculations are based on the idea to provide a lower bound on the information necessary to explain the human performance.

For example, in classical digits, cards, words and images disciplines we assume performances are perfect, even though the row grading scheme leads to raw scores which are lower than the number of correctly recalled items. The disciplines are given in rows of *k* items. A single error in a row leads to the row being counted as *k/*2, whereas with two or more errors no points are being awarded. If the last row is only recalled up to a certain item, indicating that the competitor has not memorized further, the rule is applied to the partial row.

For words and names, we use estimates based on large samples of databases. For the associative aspect in names, images and historic dates, we use a lower bound, which avoids any assumptions on cognition. We do not incorporate any of the underlying mental processes, such as the memory palace, in our calculations. We denote entropy associated to achieving a raw score *n* in a discipline by *H*(discipline, *n*).

### Names and faces

Faces with a first and a last name are provided during the memorization phase in classical competitions. In the recall period, faces are scrambled and competitors need to recall the names. A point is awarded for every correctly retrieved name, so two points are possible per face. This implies that a raw score of *n* corresponds to memorizing at least *n/*2 faces. ML national and international names feature 30 faces with only a first name.

IAM names are generated as follows:

- Names are uniformly chosen from eight different ‘regions’, which cover different language branches.
- In five-minute names, there are 12 “ long” names of eight letters or more and in the 15 min version 24 long names. The long names are chosen uniformly among first and last names.

In the memorization phase the number of names equals the world record plus 20%. As the current five-minute world record is 105, we assume a fraction 12*/*126 of long names. Katie Kermode, who co-developed the latest IAM training software^17^, provided us with a list of the numbers of short/long, male/female per region. This allowed us to calculate the entropy carried by IAM names as

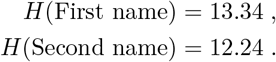

A score of *n* requires at least *n/*2 faces to be distinguished in recall. Assuming that each of the faces was instead a number, or as clearly distinguishable as possible, one could use the following strategy to solve the discipline. During memorization, one would first assemble the names in a canonical order or sort by corresponding number and then memorize the names as a list. In recall, one could recreate the order and then recall the list of names. Having a number for each face, one can see that the lower bound for the information of this strategy and thus association tasks is log_2_(#faces). Almost all of the provided information is memorized in record performances, allowing us to assume a similar amount of first and last names. We obtain the overall lower estimate

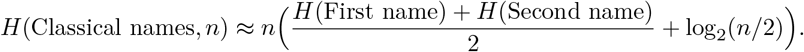

ML does not share the precise names distribution, but Simon Orton, the developer of ML, kindly applied the entropy formula for us. Both male and female names have an entropy of

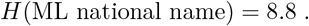

This allows us to calculate the entropy associated to a national names performance in ML as

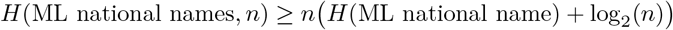

for 0 ≤*n* ≤ 30. International names consists out of all 41 national names databases combined. Simon Orton provided us again with the result of the entropy formula

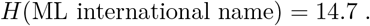

Similar to national names, we computed the formula

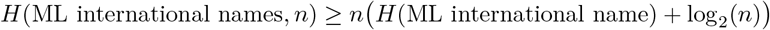

for 0 ≤ *n* ≤ 30.

### Words

Words is the only discipline for which no database for IAM competitions exists. As competitors memorize words in their native language, it is easier for organizers to provide translations for the then limited task set. As such, performances are slightly harder to compare than in other disciplines. Nevertheless, the discipline can be trained on the IAM training website^17^. We assume that players would have complained if the training mode was too easy and used a list provided by Katie Kermode. It contains 1683 concrete nouns, 829 abstract nouns and 763 verbs with some words classified for several categories. The composition of the sequence of words in competitions is 80% concrete nouns, 10% abstract nouns and 10% verbs^28^. Accounting for words being in several categories, this leads to the estimate

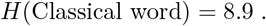

We then use the formula

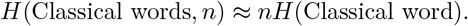

Simon Orton informed us that ML words are uniformly chosen among the ML words database. We found more than 3900 different words, giving us the lower bound

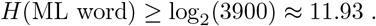

This allows us to calculate the entropy associated to a words performances in ML as

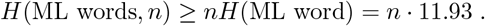

for 0 ≤ *n* ≤ 50, as 50 words are given.

### Images

In this discipline, the task is to recreate the original order of a sequence of images. While the scores reach several hundred in the classical format, it is important to note by what system the images discipline is marked. Images are presented in rows of five, and in recall each row of five images is shown in a random order and participants need to indicate the previous order by numbers. There is a deduction of one point per incorrect row preventing guessing. Enrico Maraffa, the world record holder in IAM images, confirmed in personal communication that he is not using memory palaces, but only links. Therefore, he would be able to recall the same information if the rows were scrambled. As he is able to identify the rows, similar to the associative aspect in names, we add log_2_ 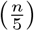 for the row identification to arrive at

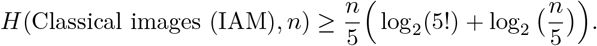

In ML, participants see 30 images and need to reassemble them during recall. Thus, the entropy of the ML images discipline is

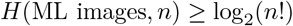

for 0 ≤ *n* ≤ 30.

### Digits

*H*(ML images, *n*) *≥* log_2_(*n*!)

The calculation of the number of bits conveyed by decimal digits and binary digits is straightforward:

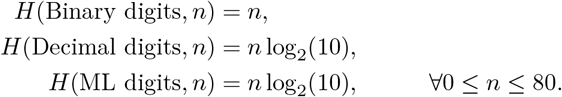

### Historic dates

The objective is to memorize the year between 1000 and 2100 of fictional historic dates. In recall, the fictional events are in a random order without years. The entropy carried by the year of a single memorized historic date is log_2_(1100) = 10.1. We again need to account for the associative component by adding log_2_(*n*). In reality encoding will be less efficient. We can thus provide a lower bound to the information processed in the discipline by

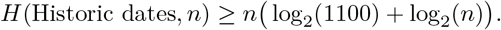

### Cards

The entropy of a subset of 52 playing cards is

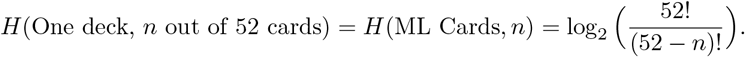

We again assumed that memorization was perfect and that any remainder mod 52 originates from the memorization of a partial deck at the end of the recalled sequence, leading to the formula

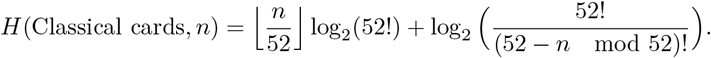

### Probabilistic model

We present a short probabilistic model relating the information-rate power law to the probability of perfect memorization in short tasks. The main idea is that competitors try to remain error-free and that a long discipline can be decomposed into shorter, independent tasks. It necessarily neglects many of the aspects described in the previous sections. We model one of the sequential disciplines. Suppose that competitors try to achieve perfect memorization to avoid incurring any of the rows being awarded no points. To do so, they aim for perfect memorization with a certain probability

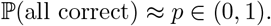

We subdivide the sequence into groups of *k* items, each with entropy *H*(item). Let *A*(*k, t*) be the event that a competitor attempts and correctly recalls *k* elements in *t* seconds memorization time excluding the reading time *r* for the group. Let *T* be the overall time of the discipline in seconds. Assuming that groups of *k* elements are independent of each other, we could then write

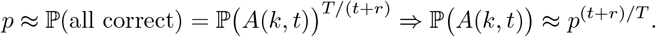

The official records suggest that the information rate scales as a power law of the discipline, so we obtain

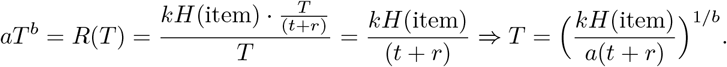

This implies that

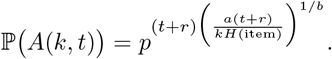

The prediction ℙ *A*(*k, t*) for fixed *k* can be found in Figure 2 **B**. If, alternatively, one fixes *t*, the model predicts that the probability to memorize all *k* items resembles a reversed logistic growth curve. For a small number *k* of items the probability is close to one. As the number of items *k* increases, the decay of the probability is first slow, accelerates and then decelerates, so that the probability asymptotically converges to zero.

## List of records

**Table 3:**
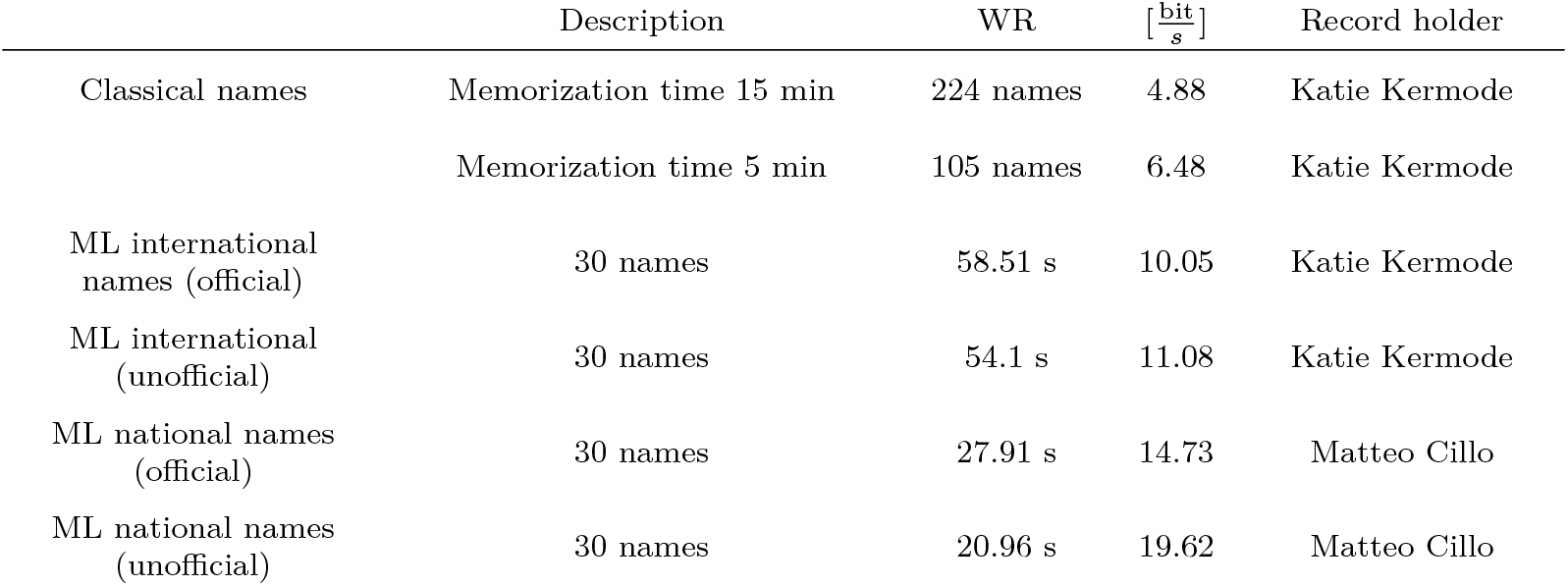
Names world records.

**Table 4:** Words world records.

| | Description | WR | $[\frac{\text{bit}}{\text{s}}]$ | Record holder |
| --- | --- | --- | --- | --- |
| Classical words | Memorization time 15 min | 318 words | 3.15 | Katie Kermode |
|  | Memorization time 5 min | 145 words | 4.3 | Yanjaa Wintersoul |
| ML words (official) | 50 words | 32.41 s | 18.4 | Don Michael Vickers |
| ML words (unofficial) | 50 words | 27.95 s | 21.34 | Don Michael Vickers |
|  | Youtube training<sup>9</sup> | 12.77 s<br>45/50 | 42.04 | Don Michael Vickers |

**Table 5:**
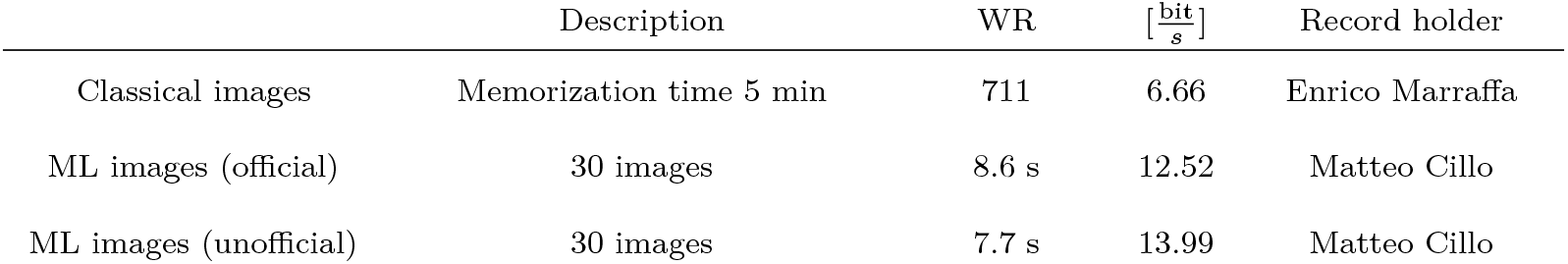
Images world records.

**Table 6:** Decimal digits world records.

| | Description | WR | $[\frac{\text{bit}}{\text{s}}]$ | Record holder |
| --- | --- | --- | --- | --- |
| Classical digits | Memorization time 60 min | 3412 | 3.15 | Orkhan Ibadov |
|  | Memorization time 30 min | 1955 | 3.61 | Sylvain Arvidieu |
|  | Memorization time 15 min | 1300 | 4.8 | Munkshur Narmandakh |
|  | Memorization time 5 min | 630 | 6.98 | Andrea Muzii |
| ML digits (official) | 80 decimal digits | 11.45 s | 23.2 | Alex Mullen |
| ML digits (unofficial) | 80 decimal digits | 10.82 s | 24.56 | Andrea Muzii |
|  | Youtube training<sup>8</sup> | 9.75 s<br>80/80 | 27.26 | Andrea Muzii |
| Speed Memory | Memorization time 4 s | 31 | 25.75 | Joaquin Garcia |
|  | Memorization time 1 s | 21 | 69.76 | Ramón Campayo |

**Table 7:** Binary digits world records.

| | Description | WR | $[\frac{\text{bit}}{\text{s}}]$ | Record holder |
| --- | --- | --- | --- | --- |
| Classical binary | Memorization time 30 min | 6270 | 3.48 | Munkshur Narmandakh |
|  | Memorization time 5 min | 1467 | 4.89 | Munkshur Narmandakh |
| Speed Memory | Memorization time 4 s | 100 | 25 | Joaquín Garcia |
|  | Memorization time 1 s | 50 | 50 | Joaquín Garcia |

**Table 8:**
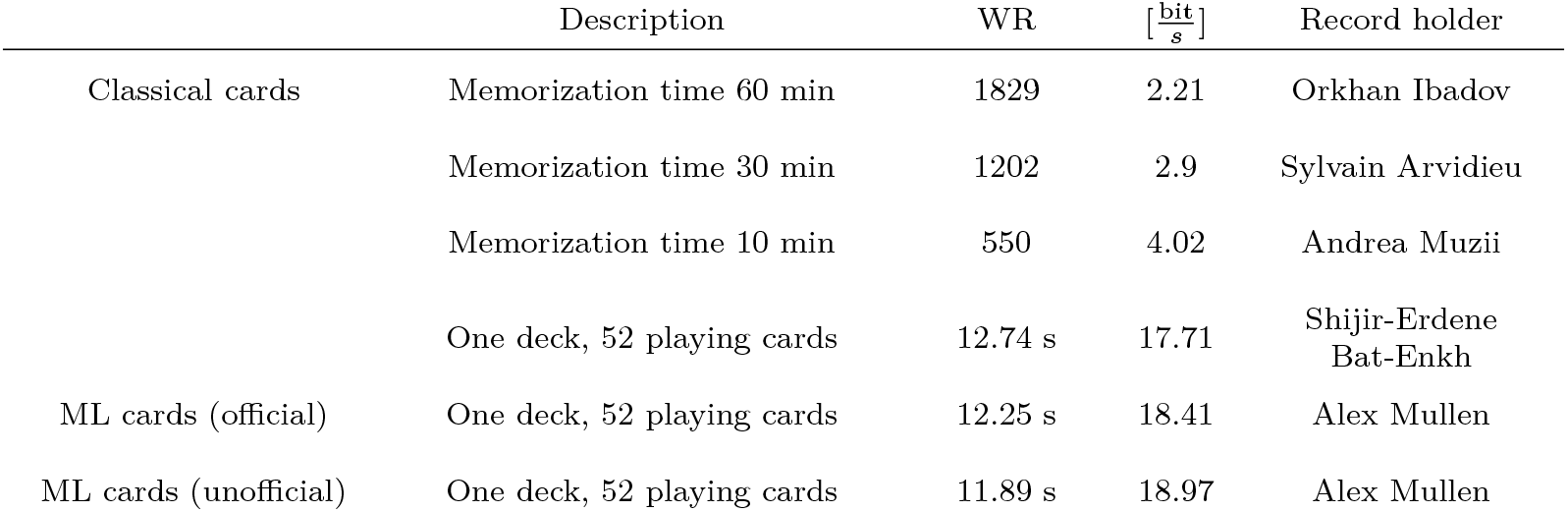
Cards world records.

